# Description of *Protosticta armageddonensis sp. nov*. (Odonata: Zygoptera: Platystictidae) from the Western Ghats of India

**DOI:** 10.1101/2023.01.11.523583

**Authors:** Arajush Payra, Reji Chandran, Ameya Deshpande, Pankaj Koparde

## Abstract

A new species of *Protosticta* Selys, 1885 (Odonata: Zygoptera: Platystictidae) is described based on two male specimens collected from Kerala, at the southern end of the Western Ghats in India. We compared *P. armageddonensis* sp. nov. with the three closely similar *Protosticta* species recently described from the Western Ghats, namely *P*.*anamalaica* Sadasivan, Nair & Samuel, 2022, *P. cyanofemora* Joshi, Subramanian, Babu & Kunte, 2020, and *P. monticola* Emiliyamma & Palot, 2016, to provide comprehensive differential diagnosis. The new species is distinguished from its congeners by a combination of characters, including the structure of prothorax, caudal appendages, genital ligula, and markings on the 8th abdominal segment. A revised key of *Protosticta* spp. of the Western Ghats, based on mature male specimens is provided.

## Introduction

The genus *Protosticta* Selys, 1885 of Family Platystictidae Kennedy, 1920; with 54 described species occurs in the South and Southeast Asian countries from Pakistan, India to Myanmar, south China, Lao PDR, Thailand, Cambodia, Malaysia, Borneo, Sulawesi and Philippines (van Tol, 2000; Paulson et al., 2022). These shade-loving, slender damselflies chiefly inhabit the undergrowth of wet shaded forests amid hill streams. The genus was established by Selys (1885) with the type species *Protosticta simplicinervis* Selys, 1885 from Sulawesi (Selys, 1885). The genus *Protosticta* can be distinguished from the closely related genus *Drepanosticta* Laidlaw, 1917 by the absence of anal bridge (van Tol, 2000). India is home to 17 species of *Protosticta*, of which 14 species (*P. anamalaica* Sadasivan, Nair & Samuel, 2022; *P. antelopoides* Fraser, 1931; *P. cyanofemora* Joshi, Subramanian, Babu&Kunte, 2020; *P. davenporti* Fraser, 1931; *P. francyi*Sadasivan, Vibhu, Nair &Palot 2022; *P. gravelyi* Laidlaw, 1915; *P. hearseyi* Fraser, 1922; *P. monticola*Emiliyamma&Palot, 2016; *P. mortoni* Fraser, 1924; *P. myristicaensis* Joshi &Kunte, 2020; *P. ponmudiensis*Kiran, Kalesh&Kunte, 2015; *P. rufostigma*Kimmins, 1958; *P. sanguinostigma* Fraser, 1922 and *P. sholai* Subramanian &Babu, 2020) are restricted to the Western Ghats Biodiversity Hotspot and the rest three (*P. damacornu* Terzani and Carletti, 1998; *P. fraseri* Kennedy, 1936 and *P. himalaica* Laidlaw, 1917) occur in the Northeastern states of India (Kalkman et al., 2020; Sadasivan et al., 2022). In the Western Ghats, most of the *Protosticta* species are confined to southern and central Western Ghats, and only a few have been reported from the northern parts of the Ghats (Rangnekar & Naik, 2014; Koparde et al., 2015; Joshi et al., 2020; Sawant et al., 2022).

Here we describe a new species of *Protosticta*, based on the two male specimens collected from Kerala, southern Western Ghats. The new species can be separated with the closely allied with *P*.*anamalaica* Sadasivan, Nair & Samuel, 2022, *P. cyanofemora* Joshi, Subramanian, Babu & Kunte, 2020, and *P. monticola* Emiliyamma & Palot, 2016, by the combination of characters, including the marking on 8th abdominal segment, structure of caudal appendages and genital ligula.

## Materials and Methods

The species was first photographed by RC on 07.xii.2022 from the type locality. Following the study of the photographic records, the specimens were captured on 27.xii.2022 using a hand-held entomological net and immediately stored in 100% ethanol. The specimens were examined and photographed using Carl Zeiss stereomicroscope (Stemi 305) with a microscope camera (MICAPS ECOCMOS510B). Field photographs were taken by a Canon 1200D digital camera. Specimens were measured with the help of a digital caliper. Systematic arrangement follows Paulson et al. (2022). Morphological terminology follows Garrison et al. (2010) and wing venation follows Riek & Kukalová-Peck (1984).

Abbreviations: Ab: anal bridge; Ca: caudal appendages; Pt: pterostigma; Ax: antenodal crossveins; Px: postnodal crossveins; S1 to S10: abdominal segments 1 to 10; FW: fore wing; HW: hind wing; RP1: radius posterior, first branch; RP2: radius posterior, second branch; IR2: intercalary vein 2; Alt: Altitude; m: meters, mm: millimeters

### *Protosticta armageddonensis* sp. nov. Chandran, Payra, Deshpande, Koparde

*Holotype*: m#, Near Merchiston Estate, (N 8.7445 N, 77.1293 E, alt: 900 masl), Thiruvananthapuram - Ponmudi Road, Thiruvananthapuram, Kerala, India, 27.xii.2022, R. Chandran, A. Payra & A. Deshpande leg.

**Paratype:** m#, location, Date of collection and collectors same as holotype.

#### Etymology

The specific name highlights the major decline in insect abundance across the globe (Hallmann et al. 2017; Leather 2017; Cardoso & Leather 2019; Cardoso et al. 2020), an event termed as Insect Apocalypse leading to the Ecological Armageddon. Though there are opposing views on the Insect Apocalypse (Goulson 2019; Schowalter et al. 2019; Montgomery et al. 2020; Saunders et al. 2020); a more cautious approach towards insect conservation is mandated in the present scenario. The concept of Insect Apocalypse is perhaps premature, but it might be a near future possibility given the current trends in insect decline across multiple groups and regions. The nomenclature is a shout out to the world leaders to help support the conservation of insect diversity on the Planet Earth through habitat conservation, long-term monitoring, and peoples’ participation in conservation. The suggested common English name is Armageddon Reedtail.

#### *Description of the Holotype Male* (Figs 1, 2 a)

**Head (Figs. 1a, 1e)**. Eyes black dorsally and posteriorly; anteriorly bluish black and pale blue below. Labium pale yellow. Labrum pale blue with black margins. Anteclypeus pale blue. Postclypeus glossy black. Frons, vertex and occiput are black with metallic sheen. Ocelli white. Base of antennae white, filaments brown.

**Figure 1.**
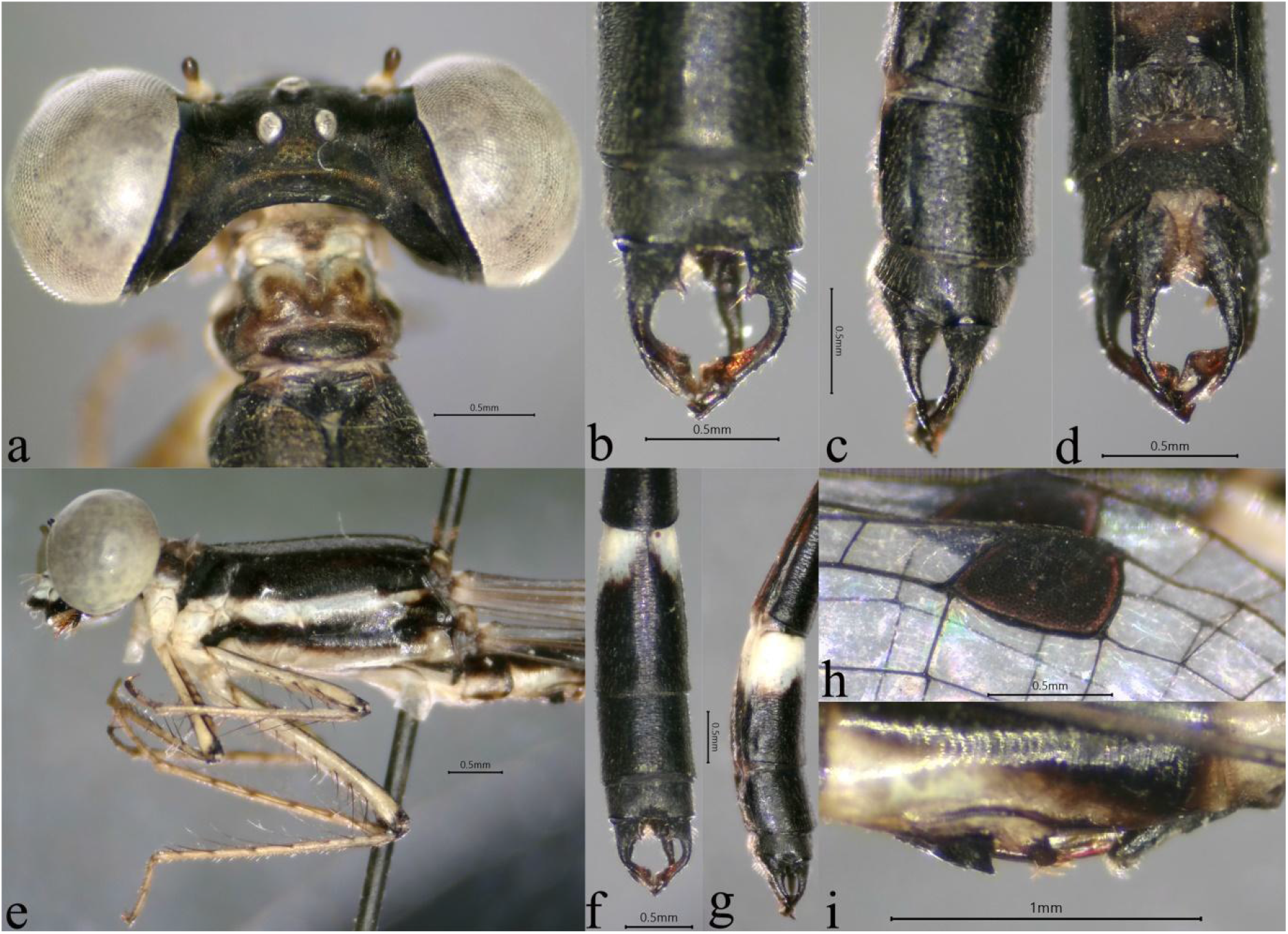
*P. armageddonensis sp. nov*. Holotype male: (a) head and prothorax, dorsal view; (b) caudal appendages, dorsal view; (c) caudal appendages, lateral view; (d) caudal appendages, ventral view; (e) head, thorax and legs, lateral view; (f) end abdominal segments, dorsal view; (g) end abdominal segments, lateral view; (h) pterostigma; (i) left lateral view of secondary genitalia.

**Figure 2.**
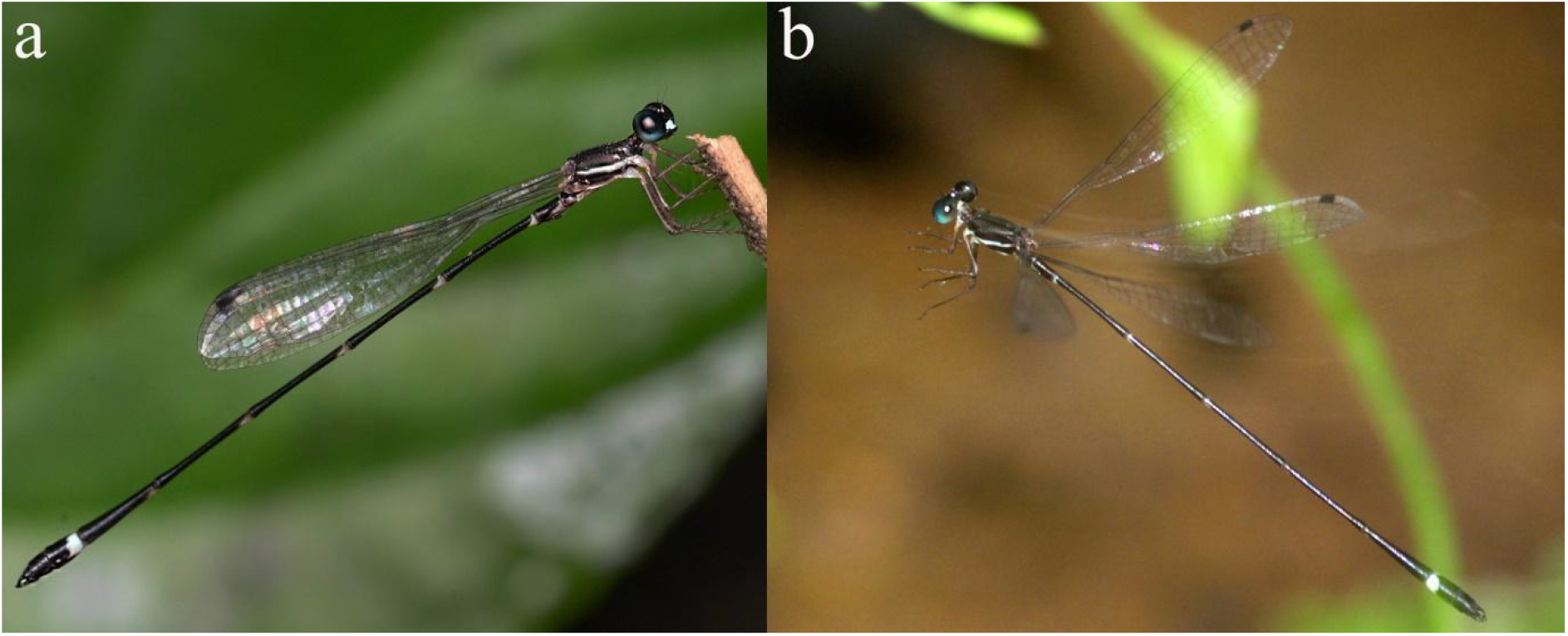
Habitus of *P. armageddonensis* sp. nov. (a) Holotype male; (b) Paratype male.

**Thorax (Figs. 1a, 1e)**. Prothorax brownish. Anterior lobe is pale brown with a dark brown spot at the center. Middle lobe dark brown with pale brown margins. Posterior lobe black dorsally and brown laterally. Dorsal half of propleuron dark brown, pale below. Synthorax black with bronze sheen. Mesepisternum and mesepimeron bronze black. Antealar triangle black. Metepisternum dark brown to black with pale blue stripes. Metepimeron is mostly pale blue; black to dark brown below the metapleural suture. Mesinfraepisterrnum black, pale towards coxa. Dorsal half of metinfraepisternum black, ventral half pale white. Legs: Coxae and trochanters white. Femora, tibia and tarsal claws are brown. Extensor surface of the femora is black.

**Wings (Fig. 1h)**. Hyaline, Pt dark brown covering one cell. Length of Pt slightly more than the breadth. Ax-2, Px-10 in all wings except left FW, where Px is 12. Ab absent. One cell between the junction of RP1-RP2 and the origin of IR2 in FW and HW.

**Abdomen (Figs. 1f, 1g, 1i, 2a)**. Black. Dorsum of S1 black with a small, dirty white crown-shaped marking at the center. S1 is white laterally, bordered with black. S2 dorsally black with a narrow elongated white sheen on the dorsal 1/3rd of the segment. S2 ventro-laterally white. White annules at the anterior border of S3-S7, which become more extensive ventro-laterally.About half the basal part of S8 marked with pale blue not connected dorsally. The pale blue marking extends ventro-laterally till the middle of the segment. S9-S10 completely black. Anterior hamuli black, large, quadrangle shaped and expanded laterally. Posterior hamulismall, oval shaped and brownish with golden yellow hairs.

**Genital ligula (Figs. 3a-3b)**. As in the figure, illustrated based on the paratype male. Distal end of the first segment with long setae..

**Figure 3.**
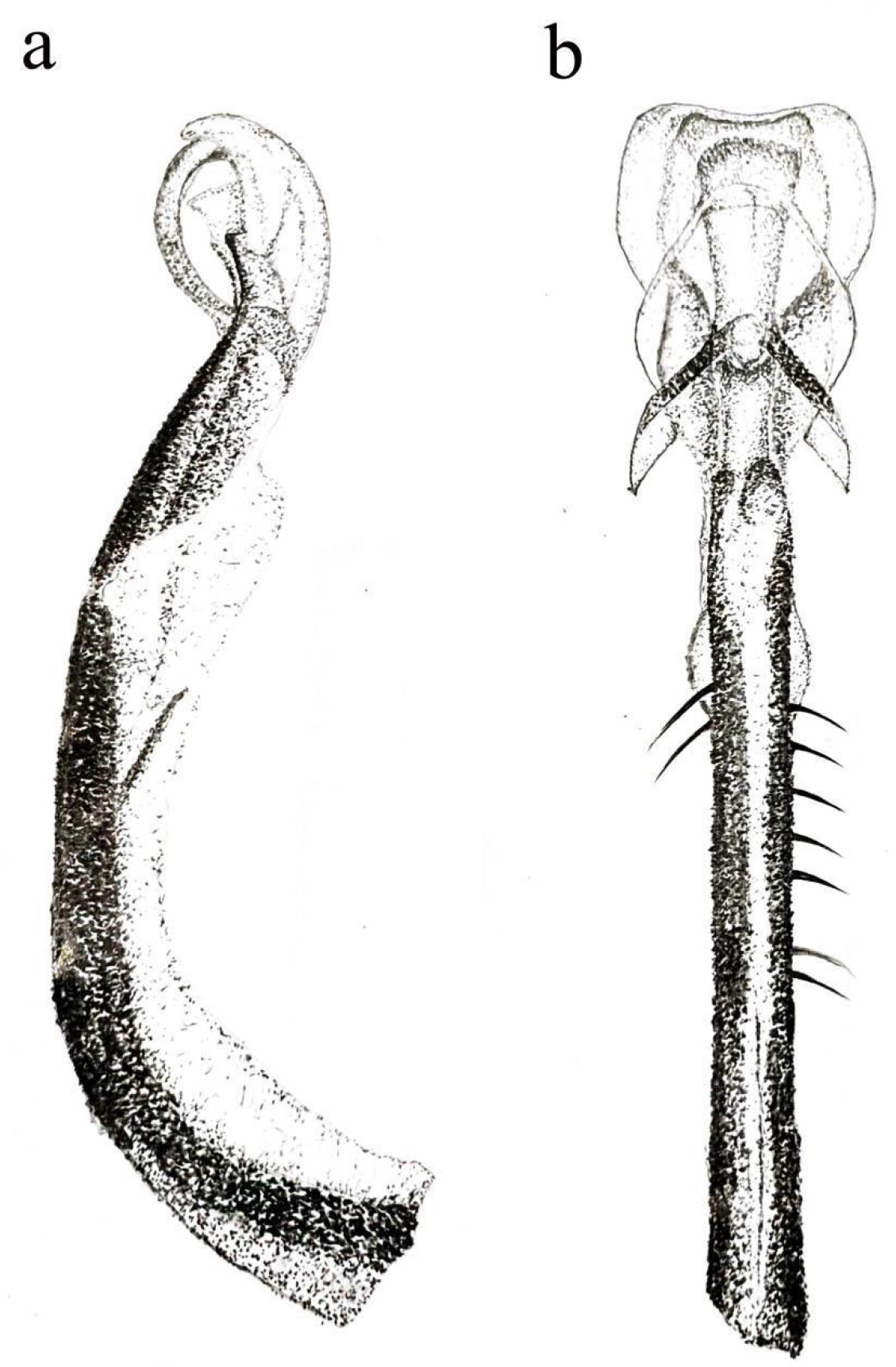
Genital ligula of *P. armageddonensis sp. nov*. (a) lateral view; (b) dorsal view

**Caudal appendages (Figs. 1b-1d)**. Black.Cerci black with brown apices, more than twice the length of S10; a basal triangular spine at the inner margin, more than 1 / 4th of the total length of the cerci. Cerci bifid at apex, outer fork thicker and curved inwardly;inner fork flattened, beak shaped and slightly shorter than the outer fork. Paraprocts black with brown apices, with a small stout spine at the inner side of the base.The basal 1/4th length of paraprocts is thicker. Apex of paraprocts pointed, dorso-ventrally flattened.

Measurements (in mm). Abdomen + caudal appendages= 29.7, FW=18, HW=17.2

### *Variations in Paratype Male* (Figs. 2b, 4)

**Figure 4.**
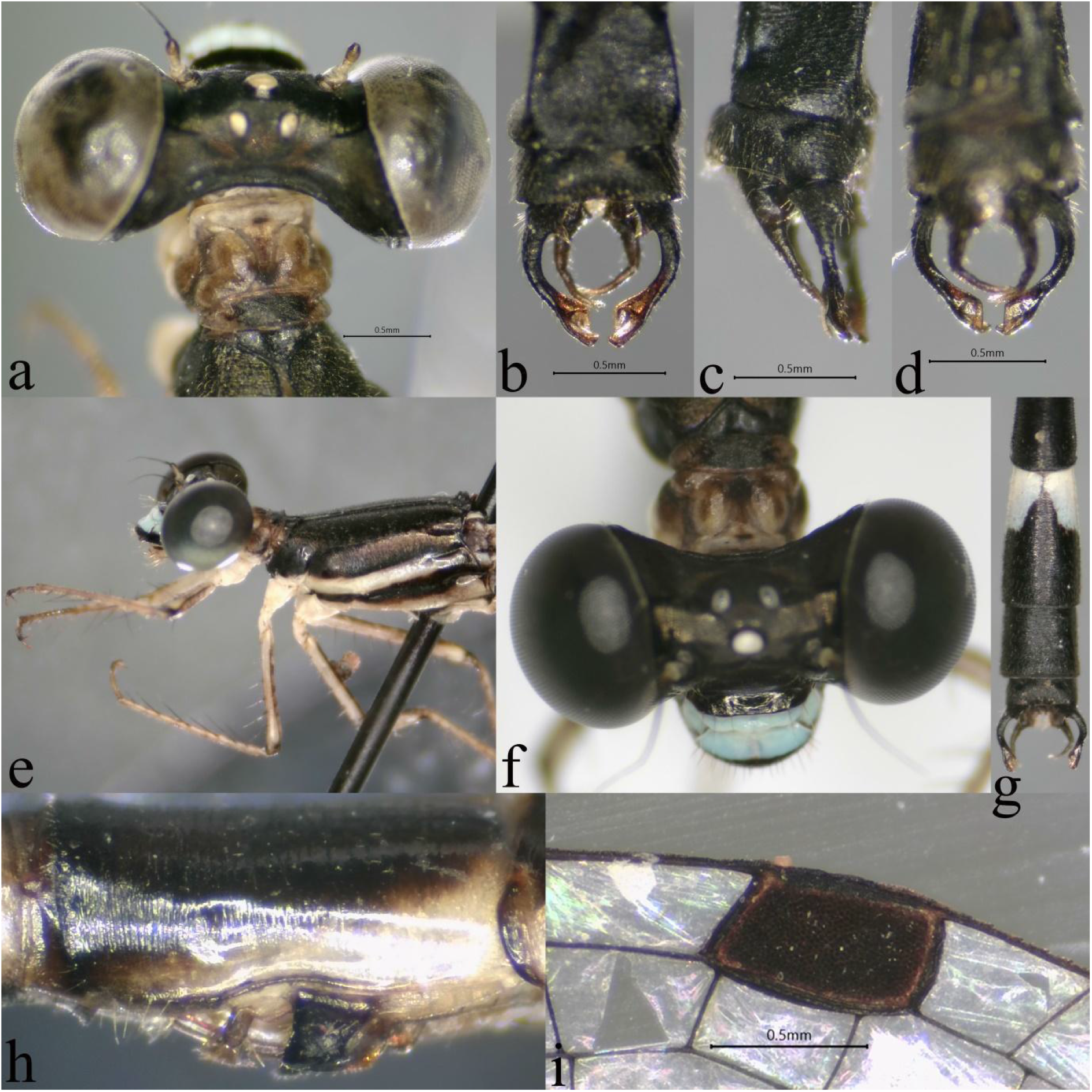
*P. armageddonensis sp. nov*. Paratype male: (a) head and prothorax, dorsal view; (b) caudal appendages, dorsal view; (c) caudal appendages, lateral view; (d) caudal appendages, ventral view; (e) head, thorax and legs, lateral view; (f) face (g) end abdominal segments, dorsal view; (h) left lateral view of secondary genitalia; (i) pterostigma.

Eyes dark brown above, bluish black anteriorly, pale blue below and laterally. Antennae brown with pale yellow scape and pedicel. Anterior lobe of the prothorax is pale brown with a small black spot at its center. Middle lobe brown. Dorsal half of propleuron dark brown, ventral half pale. Pt covering 1 and 1/5 of the cells. Px, 10 in all wings. One cell between the junction of RP1-RP2 and origin of IR2 in FW and HW except two cells in right HW. S1 dorsally black with a small white crown shaped spot at centre. Posterior hamuli are small, pale yellow, oval shaped with thick golden yellow hairs.

Measurements (in mm). Abdomen + caudal appendages= 29.5, FW=17.5, HW=17

***Female*:** Unknown

#### Diagnosis

The species is distinguished from the most *Protosticta* spp of the Western Ghats by having simple posterior lobe of prothorax lacking spines (in *P. antelopoides, P. francyi*, and *P. ponmudiensis*, posterior lobe with a pair of spines) and about anterior 1/3^rd^ of S8 pale blue colored, not connected dorsally (in *P. davenporti, P. gravelyi, P. hearseyi, P. mortoni, P. myristicaensis* and *P. rufostigma* bright blue markings on the anterior 1/3^rd^ or more of S8 dorsally connected). S9 completely black (in *P. sholai* S9 lateral 2/3^rd^ of the segment is marked yellow). Dark brown pterostigma, outer fork of cerci not bi-lobed (pterostigma red and bilobed outer fork of cerci in *P. sanguinostigma*). *Protosticta armageddonensis sp. nov*. is closely similar to newly described species *P. cyanofemora, P. anamalaica*, and *P. monticola*. In *P. armageddonensis sp. nov*., femur is brownish with black exterior, while femur is bright blue interiorly in *P. cyanofemora*, brown femur in *P. anamalaica* and pale yellow femur in *P. monticola*. S8 with pale blue annule extending laterally about ½ of the segment in *P. armageddonensis sp. nov*., but the bright blue mark extends laterally 2/3^rd^ of the S8 in *P. cyanofemora*. In *P. anamalaica*, S8 marked pale yellow ventro-laterally not reaching distal end of the segment and in *P. monticola* dorsum of S8 marked black, ventro-laterally yellow extends to its distal end. Also, the new species can be separated from *P. cyanofemora, P. anamalaica* and *P. monticola* by its smaller size, eye coloartion, and size of setae (Table 1).

**Table 1.**
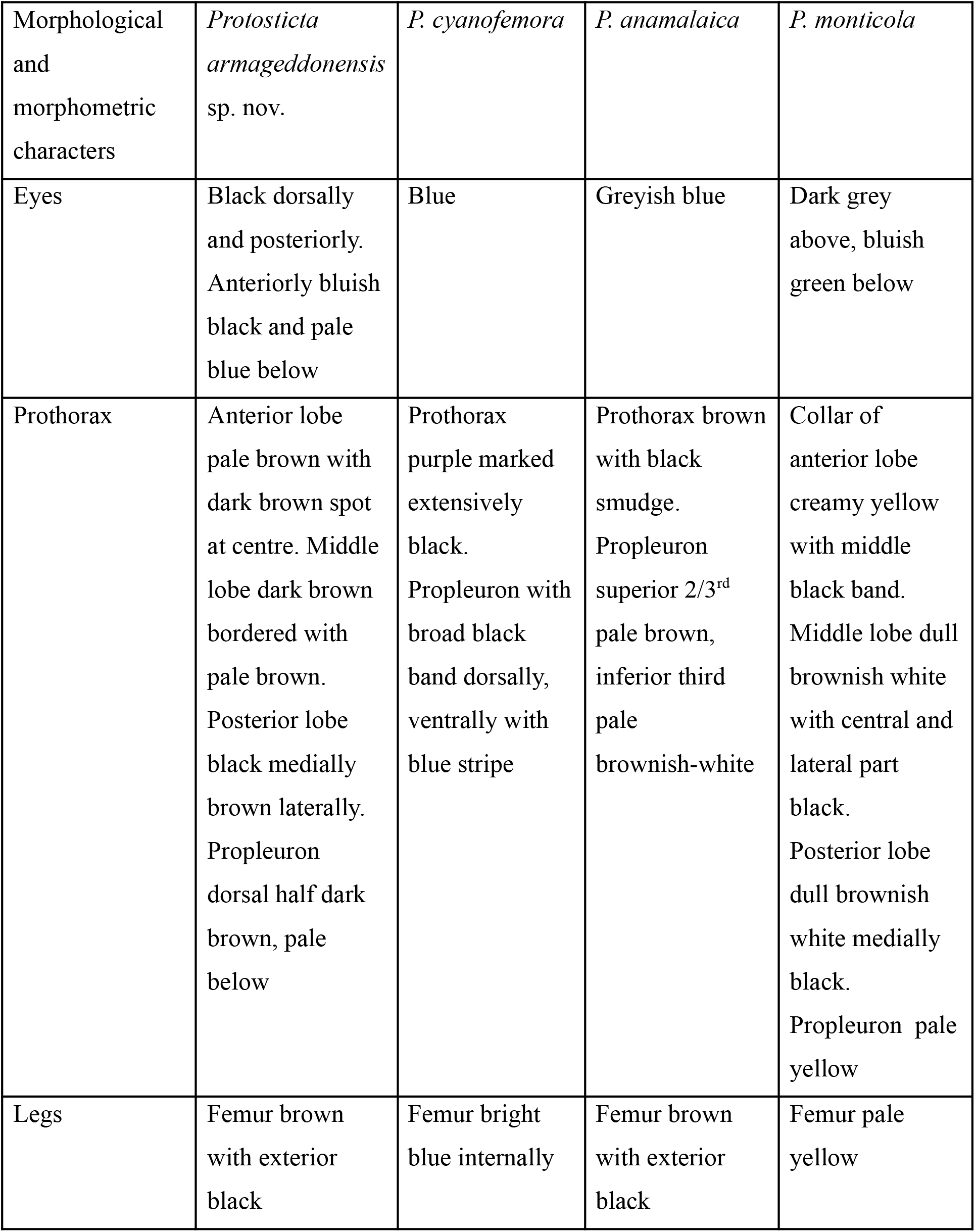
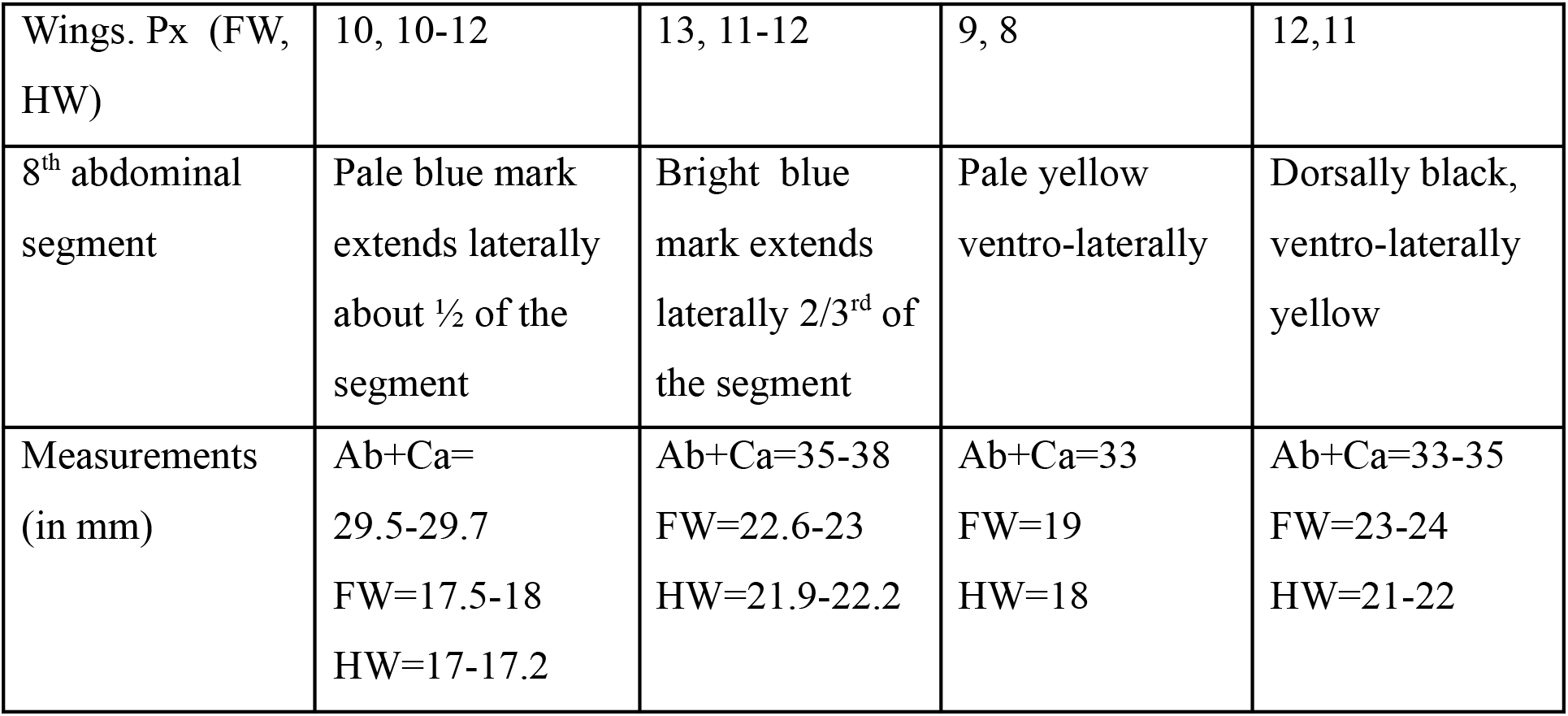
Comparison of morphological and morphometric characters based on mature male specimens of *Protosticta armageddonensis sp. nov*., *P. cyanofemora, P. anamalaica* and *P. monticola* based on Emiliyamma & Palot (2016), Joshi et al. (2020) and Sadasivan et al. (2022).

### Habitat & Ecology

The species was observed along the Thiruvananthapuram-Ponmudi Road, near Merchiston Estate, Thiruvananthapuram, Kerala, at an elevation of 900 masl. Both individuals were seen perched on shrubs very close to the ground (about a foot above the ground) near a brook, in the shade of an evergreen canopy. Other odonate species observed in the same habitat were *Esme mudiensis, Protostcita gravelyi, P. ponmudiensis*, and *Hylaeothemis apicalis*.

## Acknowledgements

The work was funded by the Department of Science and Technology, Government of India (DST-SERB/SRG/2020/000190). We acknowledge efforts by Robia Kshetrimulayam for helping with illustrations. We thank Vivek Chandran for reviewing the draft manuscript. We thank David Raju for guiding us on the field. We are thankful to the Kerala State Forests & Wildlife Department for research permission (KFDHQ-4231).

## Key to the *Protosticta* spp. of the Western Ghats based on mature males modified from Joshi et al. (2020), Sadasivan et al. (2022) and Vijayakumaran et al. (2022)

1. The posterior lobe of prothorax with a pair of long spinesspines…………………………………………….2
  - The posterior lobe of prothorax without spines……………………………………………………………..4
2. The posterior lobe of prothorax with very long medial spines, extending posteriorly to a distance more than twice the length of the mesostigmal platemesostigmal plate………………………..*P. antelopoides*
  - Posterior lobe of prothorax with short medial spines extending posteriorly till the apex of the mesostigmal plate or shortershorter…………………………………………………………………………………….3
3. Lateral spines on prothorax well developed, short triangular with tip directed anterolaterally; medial spines short, never extending posteriorly beyond the apex of the mesostigmal plate; tip of cerci broad at base and tapering distallydistally………………..*P. ponmudiensis*
  - Lateral spines on prothorax rudimentary, reduced to an angular projection on the prothoracic collar, and its tip directed laterally; medial prothoracic spine long, thin, extending posteriorly just beyond the apex of the mesostigmal plate; tip of cerci narrow at base and expanding distally…………………………………………………………………………………*P. francyi*
4. Anterior 1/3rd or more of S8 bright turquoise blue connected dorsally………………………….5
  - Anterior 1/3rd of S8 yellow or blue, not connected dorsally………………………………………….10
5. Apical fork of cerci deeply incised more than 1/3rd of the total length…………………………..6
  - Apical fork of cerci shallow incised, much less than 1/3rd of total length ……………………….8
6. Cerci with a small tubercle at middle of the apical fork; length of abdomen + caudal appendages < 25 mm…………………………………………………………………………….*P. myristicaensis*
  - Cerci without such a tubercle at its center; length of abdomen + caudal appendages >25 mm. mm……………………………………………………………………………………………………………………………7
7. Prothorax with a hexagonal black marking covering central portion of posterior lobe and a small portion of the middle lobe; cerci with a prominent laterally pointed basal spine; the paraprocts with an inner stout spine at base…………………………………………*P. gravelyi*
  - Anterior & middle lobes of prothorax colored blue, no hexagonal black mark; cerci with a small laterally pointed basal spine; paraprocts without an inner spine at base………..*P. mortoni*
8. Prothorax completely blue; length of abdomen +caudal appendages <30 mm; inferior lobe of cerci very short, superior lobe not expanded………………………………………..*P. hearseyi*
  - Anterior and middle lobes of prothorax pale yellow, posterior lobe partially or completely black; length of abdomen + caudal appendages > 30 mm; inferior lobe of cerci more than 1/3^rd^ length of superior lobe, the latter expanded………………………………………………….…9
9. Dorsum of middle portion of posterior lobe of prothorax completely black extending as two points to the dorsum of middle lobe; inner fork of cerci thin and small, superior lobe rounded at apices and more than twice the length of inferior………………………..…*P. davenporti*
  - Dorsum of posterior lobe of prothorax black, laterally brown; middle lobe of prothorax with a small dorsal faint black spot; inner fork of cerci thick, superior lobe ending in a quadrangle, less than twice the length of inferiornferior…………………………………………………*P. rufostigma*
10. S9 completely black or marked only at ventral border; posterior border of prothorax not expanded; paraprocts not clubbed at apices……………………….…………………………………11
  - S9 laterally marked with a large yellow spot at anterior border, reaching more than 2/3rd of the segment, not connected apically in both sexes; posterior border of prothorax expanded; paraprocts thin, long and clubbed at apices………………………………………………………….*P. sholai*
11. Pt red; cerci with a prominent and robust basal spine; tip of the superior lobe of cerci bilobed……………………………………………………………………………………………….*P. sanguinostigma*
  - Pt black or brown; cerci with a small blunt basal protuberance, inwardly pointed; tip of the outer fork of cerci not bilobed…………………………………………………………………………………….12
12. S8 with pale blue or bright blue annule extends laterally ½ to ⅔ rd of the segment……13
  - S8 with yellow or pale yellow annule extends ventro-laterally about the entire segment….14
13. Eyes black above and blue below; femur brown with extensor surface black; S8 with pale blue annule extended laterally ½ of its length; S8 annule narrowly separated on dorsum; outer margin of cerci sinuous, tip of the superior lobe of cerci bent inwards at apices .……………..….. ***P. armageddonensis* sp. nov**.
  - Eyes blue; femur bright blue internally; S8 with bright blue annule extended laterally 2/3^rd^ of its length; S8 annule widely separated on dorsum; outer margin of cerci including superior lobe comparatively straighter in dorsal view………………………………………………………………………….…*P. cyanofemora*
14. Femur pale yellow internally; S8 black dorsally and ventrally yellow extends to its distal end; tip of the superior lobe of cerci bent inwards……………………………….*P*.*monticola*
  - Femur brown with black extensor;S8 with pale yellow annule extending ventro-laterally not reaching distal end; tip of the superior lobe of the cerci straight in dorsal view………………………………………………………………………………*P. anamalaica*

